# PEPA test: fast and powerful differential analysis from relative quantitative proteomics data using shared peptides

**DOI:** 10.1101/158212

**Authors:** Laurent Jacob, Florence Combes, Thomas Burger

## Abstract

We propose a new hypothesis test for the differential abundance of proteins in mass-spectrometry based relative quantification. An important feature of this type of high-throughput analyses is that it involves an enzymatic digestion of the sample proteins into peptides prior to identification and quantification. Due to numerous homology sequences, different proteins can lead to peptides with identical amino acid chains, so that their parent protein is ambiguous. These so-called shared peptides make the protein-level statistical analysis a challenge, so that they are often not accounted for. In this article, we use a linear model describing peptide-protein relationships to build a likelihood ratio test of differential abundance for proteins. We show that the likelihood ratio statistic can be computed in linear time with the number of peptides. We also provide the asymptotic null distribution of a regularized version of our statistic. Experiments on both real and simulated datasets show that our procedures outperforms state-of-the-art methods. The procedures are available via the `pepa.test` function of the `DAPAR` Bioconductor R package.

## 1 Introduction

Quantitative proteomics refers to the identification and quantification of the proteins present in a biological sample. This field has rapidly grown mature over the last decade, allowing for a refined understanding of a wide variety of biomolecular processes: phenotypes of new forms of life, such as giant viruses (Philippe *and others*, 2013), host-pathogen cell interactions (Le Roux *and others*, 2015) or microbial infections (Hodille and *others*, 2016). As many other omics sciences, it is based on a large scale sequencing approach, that is bound to high throughput measurements whose statistical processing is a central issue. The most classically used measurement pipeline (Commentary, 2014) is referred to as *relative bottom-up MS/MS quantification* (Zhang *and others*, 2013).

The term *bottom-up* refers to the fact that proteins are not directly identified: instead, they are first digested by an enzyme into smaller molecules called *peptides*, that are easier to analyze by MS/MS. The term *MS/MS* refers to the fact that two kinds of mass spectrometry (MS) measurements are alternatively performed. The first one is used to estimate the mass of all the peptides that are simultaneously co-analyzed and deduce their abundances. The second one garners evidence on the chemical composition of the few most abundant ones, leading to their identification. However, the abundance measure given by the first MS-measurement is influenced by many physico-chemical parameters, making it difficult to precisely relate it to a concentration measure. One therefore relies on isotope labeling to infer the real concentrations, or limits oneself to relative quantification, where only ratios are accounted for. As the MS-response of a given peptide is roughly linear in its concentration in a given chemical environment, its intensity ratio between two samples of a same experiment is well-correlated to its concentration ratio. This instrumental pipeline is particularly useful in discovery proteomics, where the goal is to find a shortlist of proteins that are significantly differently abundant. Several samples are collected under different biological conditions (*e.g.*, in healthy vs. disease, wild-type vs. mutant, etc.) and analyzed with the aforementioned pipeline, leading to a list of identified peptides and their intensity for each sample.

Deciding which proteins are differentially abundant from peptide level observations is made difficult by the presence of *shared peptides* (as opposed to *protein-specific* ones): due to the numerous homology sequences between different genes, some peptides can belong to several distinct proteins. This problem has long been reported in the literature (Jin *and others*, 2007) and according to Dost *and others* (2012), up to 50% of peptides can be shared in the proteome of complex organisms. However, to the best of our knowledge the few solutions available in the literature have hardly spread to proteomics platforms, as they are hampered by practical issues. In this context, our contribution is four fold:

1. We introduce a linear normal model which relates measured peptide MS intensities to latent protein abundances. This model can be used to build a PEptide based Protein differential Abundance (PEPA) likelihood ratio test which accounts for both protein-specific and shared peptides.
2. The resulting model involves an *nq* × (*p* + *q*) design matrix, where n is the number of samples, *q* the number of peptides and *p* the number of proteins, which makes estimation intractable in practice using naive least square implementations. We show that our likelihood ratio statistic can be computed in 𝒪(*nq*) nevertheless, making it compatible with proteomic platform throughput.
3. We empirically observe that regularized estimators of the variance parameter lead to more powerful tests. We show that under the null hypothesis of homogeneity, the regularized log-likelihood ratio statistic is still asymptotically χ^2^ distributed up to some normalization.
4. We provide R code for all our methods in the DAPAR Bioconductor R package, so they can be routinely used by proteomic practitioners.

## 2 State-of-the-art

Leaving aside the various preprocessing steps that are necessary to account for *e.g.* batch effects or missing values (see for instance Wieczorek *and others* (2016)), methods for differential analysis of proteomic datasets can be divided in two main families: *peptide-based* and *aggregation-based* methods, also referred to as summarization-based in Goeminne *and others* (2015). In the latter ones, peptide-level information is first aggregated at the protein level and proteins are then tested for differential abundance using these summaries. Peptide-based models on the other hand do not rely on an aggregation step and build a test statistic using peptide intensities as a sampling unit.

### 2.1 Aggregation-based models

Although both families allow to rank proteins according to their significance, aggregation-based methods are much more widely used on proteomics platforms than peptide-based models, as protein level abundance values make more sense to many practitioners and are easier to interpret.

The most commonly used aggregation methods avoid the issue of shared peptides by only considering protein-specific ones: either all of them, so as to involve as much information as possible in the process, or only the most abundant ones (Silva *and others*, 2006), which are best identified and least error-prone. Alternatively, all peptides can be retained for each protein whether they are shared or not. The intensities of the retained peptides are then summed or averaged, sometimes after being weighted by protein-level information such as the coverage of the protein by MS-observable peptides (Schwanhausser *and others*, 2011). We compare these approaches in Section A of the supplementary material, and show that summing or averaging all protein-specific peptides provides the best results.

More sophisticated aggregation methods have been proposed to account as precisely as possible for shared peptides. Dost *and others* (2012) split their intensities between their parent proteins by recasting the problem into a resource allocation framework, for which efficient optimization techniques are available. To the best of our knowledge (as the precise algorithm is not published and the code no longer available), it provides best results when an MS-detectability coefficient for each peptide is specified. In practice, such coefficients are scarcely known, limiting the applicability of the method to routinely analyzed and well-characterized proteomes. SCAMPI (Gerster *and others*, 2014) relies on a linear model that accounts for peptide-protein relationship like our method. However, it was designed to quantify protein abundances in single sample experiments by means of isotope labelling, and it does not generalize to joint estimation across several samples or hypothesis testing, as precisely illustrated in Section B of supplementary material.

After the aggregation step, a test for differential abundance is performed at the protein level. The most widely used procedure is the Student t-test, as well as its regularized versions SAM (Tusher *and others*, 2001) and Limma (Smyth, 2005). We present an empirical study of these different test statistics in the context of protein differential abundance analysis in Section C of supplementary material.

### 2.2 Peptide-based models

Despite the more general usage of aggregation-based methods, it has long been proposed to directly work at the peptide level, using a regression model on all the peptide intensities that corresponds to a same protein. Goeminne *and others* (2015) suggest that these approaches yield better performances than aggregation-based methods. To the best of our knowledge, the first wide-spread implementation of such a method is MSstats (Choi *and others*, 2014), on the basis of preliminary works from the same group (Clough and *others*, 2009, 2012). More recently, Goeminne *and others* (2016) proposed another implementation including regularized estimators and a robust loss function.

Most peptide-based models discard shared peptides and apply the regression only to the protein-specific peptides. A few of them however attempt to account for shared peptides, yet none of them seem amenable to statistical inference in our context where hundreds of proteins are involved. Bukhman *and others* (2008) exploit a model akin to the one we use in this paper but include a factor representing the peptide-specific relationship between the measured peptide intensity and its actual abundance. This factor causes the negative log-likelihood to be non-convex in the set of parameters making its minimization non-trivial and possibly expensive - the authors restrict themselves to peptides shared by no more than two proteins. No algorithm or code is available for this method, to the best of our knowledge. More recently Blein-Nicolas *and others* (2012) have proposed AllP, which is also based on a similar model as our work but uses a log-normal model. The use of this distribution corresponds to a common assumption on observed peptide distribution (Podwojski *and others*, 2010). Unfortunately, maximizing the corresponding likelihood is more computationally demanding than that of the normal distribution we use, as no closed form maximizer is available. As reported in its original article, a synthetic datasets with 100 proteins requires 3 days of computation and for such a dataset the algorithm does not converge in 18% of the cases. As a result, it was impossible for us to apply AllP on the much larger (real or simulated) datasets that are considered in this work.

## 3 Methods

We consider a proteomic experiment measuring the intensity of q previously identified peptides that map onto a set of p known proteins. A biological sample consists of observed intensities for q peptides. Each peptide in turn can belong to several among p proteins, and the abundance of a peptide is the sum of the abundance of all proteins containing this peptide. Formally, if the proteins have respective abundance values *θ*_1_,…,*θ*_*p*_ in a sample, then the abundance of peptide k in this sample should be 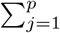, where *x_kj_* = 1 if peptide *k* belongs to protein *j*, 0 otherwise.

### 3.1 Model

The observed intensities 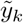 from an MS/MS experiment are typically modeled as samples from a log-normal distribution (Bukhman *and others*, 2008; Podwojski *and others*, 2010; Blein-Nicolas *and others*, 2012):

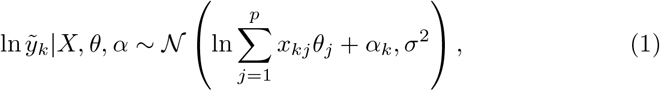

where σ^2^ > 0 is the variance of the distribution, α_*k*_ is a peptide-specific effect, X ∈ {0,1}^*q×p*^ is a binary matrix whose elements are the *x_kj_* and *θ* ∈ ℝ^*p*^ and α ∈ ℝ^*q*^ are vectors containing the protein abundances and peptide effects respectively.

The parameters of interest for differential analysis are the protein abundances *θ*_1_,…,*θ*_*p*_. They are unobserved, and we want to test whether they change between two experimental conditions of interest. More precisely, we assume the n = *n*_1_ + *n*_2_ biological samples are measured under two different experimental conditions (*n*_1_ under the first condition, *n*_2_ under the second) and we want to test

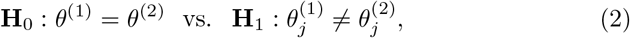

where *θ*_(1)_ and *θ*_(2)_ ∈ ℝ^*p*^ are protein abundance vectors under the two conditions and *j* is the protein being tested for differential abundance.

The data we use to test (2) consist of *q* × *n* i.i.d. intensity measurements 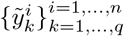. To make the analysis and computation easier, we make the approximation that:

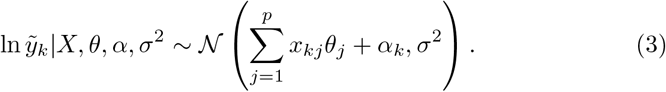

(3) makes less sense than (1) as a data generating model and loses the interpretation of *θ* as protein abundances. Nevertheless, it leads to easier inference on *θ* and we found it to yield good empirical performances, even on real data or simulated ones from log-normal distributions. In the rest of this paper, we therefore assume that the 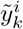 are sampled from (3) and let 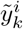 denote the log intensities ln 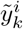.

The maximum likelihood (ML) estimators of β = (*θ*, α) for observations 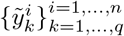 from (3) under **H**_0_ and **H**_1_ are obtained by ordinary least square:

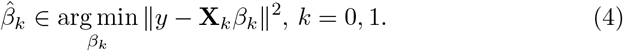

**X**_0_ is a vertical concatenation of *n* copies of the ℝ^*q*×*p*+*q*^ (*X I_q_*) matrix:

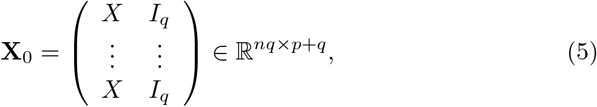

where *I_q_* is the identity matrix in ℝ^*q*^, and **X**_1_ is a vertical concatenation of n_1_ copies of the ℝ^*q*×*p*+*q*+1^ (*X*_-j_ *X*_j_ 0 *I_q_*) matrix and *n*_2_ copies of the ℝ^*q*×*p*+*q*+1^ (*X_j* 0 *X*_j_ *I_q_*) matrix: 

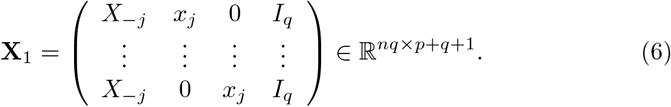

*X__j_* is the *X* matrix without its *j*-th columns *X_j_*. The matrix *y* ∈ ℝ^*nq*^ contains all 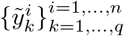, *i.e.*, the *n*_1_ peptide intensity measurements under the first condition, followed by *n*_2_ ones under the second. Finally, β_0_ = (*θ*, α) ∈ ℝ^*p*+*q*^ and 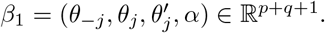.

Considering σ as a fixed parameter, the ML estimator of σ^2^ is

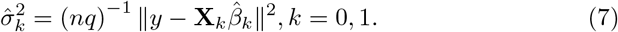

Using an inverse gamma prior on the variance σ^2^ ∼ Inv-Gamma(-1,β), the maximum a posteriori (MAP) estimator of σ^2^ becomes

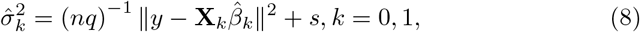

with *s* = 2β*n*^-1^. This estimator is implicitly used in test statistics like SAM (Tusher *and others*, 2001). It amounts to regularizing the variance estimate and can lead to better power than t-tests to detect differential abundance when only few samples are available. To choose s in practice, we generalize the heuristic of (Tusher *and others*, 2001): we compute our statistic for all proteins across a grid of values of s and retain the one leading to the smallest coefficient of variation of the statistic across variance levels. The motivation of the heuristic is that the amplitude of the regularized statistic should not be determined by the variance of the residuals.

Individual effects such as our peptide effect α_*k*_ are commonly modeled as random variables and endowed with a prior distribution. In our experiments, using a fixed or random α_*k*_ made little difference so we opted for the fixed effect model, as it is amenable to fast computation using Proposition 1 (see below), whereas inference under the mixed model becomes intractable as soon as a large number of peptides is shared across a set of proteins.

Finally, in practice *X* often involves disjoint sets of proteins with no peptide in common. When testing (2) for a protein j, we only use peptides whose observation affects the estimation of *θ*_*j*_. Concretely, we identify connected components in the bipartite graph whose nodes are peptides and proteins with edges between each peptide and its parent proteins, and apply our procedure to each connected component separately.

### 3.2 Computation of the test statistics

We consider the likelihood ratio statistic for (2) under model (3):

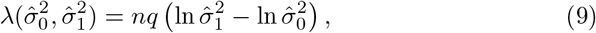

where 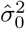 and 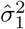 are obtained by solving either (7) for ML estimation or (8) for MAP estimation. Both estimators require solving the least square problem in (4) for *k* = 0 and *k* = 1. A naive implementation explicitly storing the *nq* × (*p* + *q*) and *nq* × (*p* + *q* + 1) design matrices **X**_*k*_ would fail when dealing with thousands of peptides even for small *n* (this was the case for the lm function of R in our experiments). Solutions of (4) can be obtained from 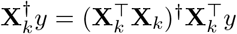 where **X**^†^ denotes the pseudo-inverse of **X**. Both 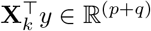 and 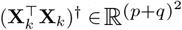 are amenable to computation but still take several minutes for each protein, which is impractical when working with large sets of proteins.

Using the fact that (9) only requires to know the sum of the squared residuals min_β_ ||*y* — **X**_k_β||^2^ and not necessarily 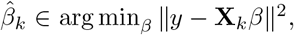, we now show how it can be computed in linear time with the size of *y*.

#### Proposition 1.

*Let* **X**_0_ *and* **X**_1_ *be defined as in* (5) *and* (6) *respectively and y* ∈ ℝ^*nq*^ *contain the n stacked* 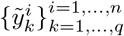 *samples, then*

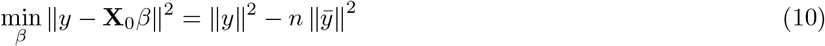

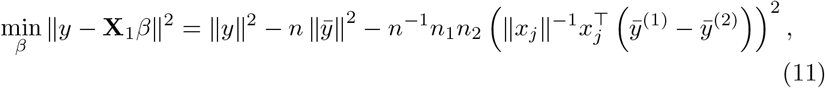

where 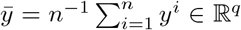 *is the average across the n samples and* 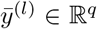 *is the average across the n_l_ samples under condition l* ∈ {1, 2}.

*Proof*. The residuals can be re-writen using the Pythagorean theorem: 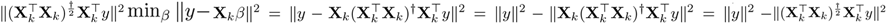, where 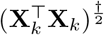 denotes the square root of the pseudo-inverse of 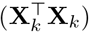.

For 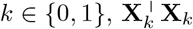 admits the Cholesky decomposition

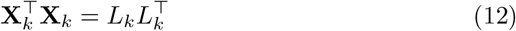

with

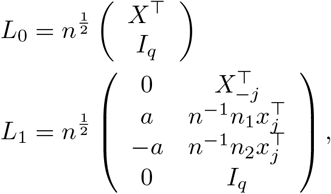

where *a* = *n*^-1^ (*n*_1_*n*_2_)^1/2^ ||*x_j_*|| We also notice that

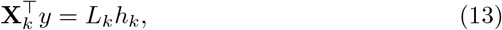

with

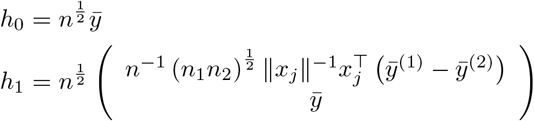

Combining (12) and (13), we obtain 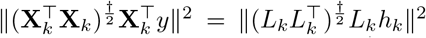. Since *L*_0_ has full column rank *q* and *L*_1_ has full column rank q+1, 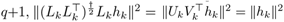, where 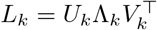 is the singular value decomposition of *LL_k_*.

When *X* is a binary matrix, 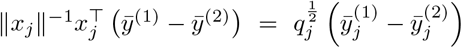 where 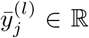, *l* = 1, 2 is the average across samples under condition l of the log-intensities of all peptides belonging to protein *j* and *q_j_* = ||*x_j_*||^2^ is the number of peptides belonging to protein *j*. The additional term in 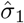 can then be interpreted as the squared difference between the average log intensities across peptides between the two conditions.

An important consequence of Proposition 1 is that the log-likelihood ratio statistic (9) can be computed in 𝒪(*nq*) by computing averages of subsamples of y, without storing the **X**_*k*_ matrices or diagonalizing the 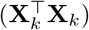 matrices.

### 3.3 Null distribution of λ

By Wilk’s theorem (Wilks, 1938), we know that λ converges in law to a 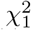 distribution as *nq* → ∞ and the *y_ik_* are sampled i.i.d. under **H**_0_ (*i.e.*, *θ*^(1)^ = *θ*^(2)^) and when using maximum likelihood estimators of (*θ*, α, σ^2^) from (4) and (7). This result provides asymptotic levels for our test, as rejecting *H*_0_ when 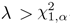, where 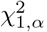 is the 1 — α quantile of the 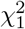 distribution, asymptotically leads to a false positive rate of α. When using a MAP estimator (8) for σ^2^, Wilk’s theorem does not hold anymore, and indeed we observed in our experiments that the null distribution of λ under *H*_0_ deviates from the 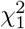 distribution. However, Proposition 2 shows that multiplying λ by a constant factor is enough to recover a correct asymptotic level.

#### Proposition 2.

*Let* 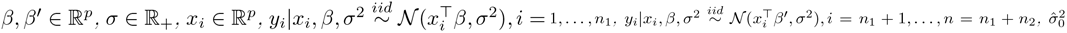 *and* 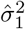 *denote the maximum likelihood estimator of* σ^2^ *under* **H**_0_: β = β’ *and* 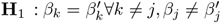 *respectively, s* ≥ 0 *and* 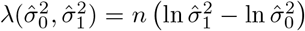. *If β = β’, then*

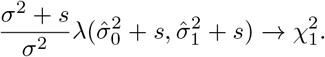

#### Proof.

We first derive the null distribution of 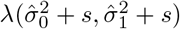 when

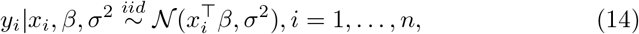

and 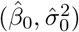 and 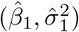 are the maximum likelihood estimators of (β, σ^2^) under **H**_0_: β_j_ = 0 and **H**_1_: β_j_ =0 respectively.

We follow the line of proof of Wilk’s theorem in (van der Vaart, 2007, Section 16.2). Classical asymptotic normality of the maximum likihood estimator provides that

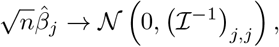

where

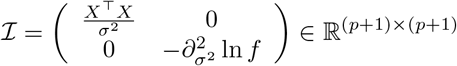

is the Fisher information matrix with respect to parameters (β, σ^2^) for the normal distribution *f* described in (14), and *X* ∈ ℝ^n×p^ is the matrix whose rows are the *x_i_*.

We note that

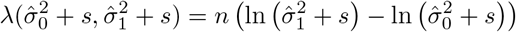

can also be written as 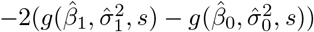

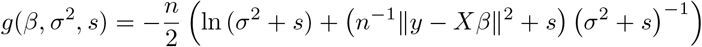

where

Even though *g*(β, σ^2^, *s*) is not the log-likelihood of the *y_i_*, its gradient 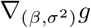 is cancelled at the maximum likelihood estimator 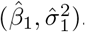. Using a second order Taylor approximation of 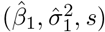 around 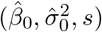, one can therefore show that

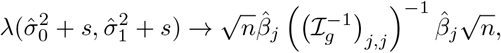

Where *I_g_* is minus the expectation of the Hessian matrix of *g*:

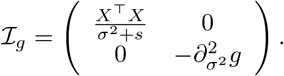

It follows that λ → *z*_2_, where *z* ∼ 𝒩(0,σ) and

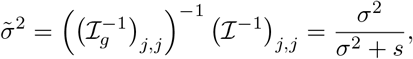

and therefore 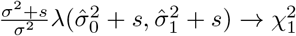.

The proposition follows using a change of variable:

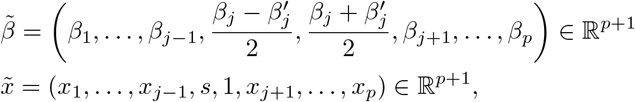

where *s* is 1 for the first *n*i *x_i_* and —1 for the remaining *n*_2_.

Alternatively, it was brought to our attention that this result could be derived as a special case of Theorem 1 in Harchaoui *and others* (2008), which describes the distribution under the null hypothesis of homogeneity for a statistic built over samples in arbitrary spaces. Their statistic is a kernel version of the Hotelling *T*^2^ statistic, boiling down to a squared t-statistic in the unidimensional case which corresponds to the likelihood ratio statistic for our model. In the univariate case, their general distribution also boils down to a 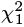 distribution.

In practice, σ^2^ is unknown but we obtained satisfying test levels on both real and simulated data by using the maximum likelihood 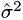 instead.

## 4 Experimental setting

We use both simulated and real data to evaluate the performance of our test procedure. Simulated datasets allow us to control key parameters such as the proportion of shared peptides, but the conclusions we draw only hold for data which behave like the simulations. On the other hand, because of the difficulties that are inherent to the wet-lab procedures (see Section 4.2), it is not possible to prepare real samples with an indisputable ground truth and which contain an important proportion of shared peptides. As a result, real and simulated datasets provide complementary views on the performance evaluation.

### 4.1 Simulated datasets

We simulate peptide intensities for each of *n*_1_ = *n*_2_ = 3 samples under two biological conditions, for *q* = 5000 peptides belonging to *p* = 1000 proteins. We purposely use a generative model which differs from the normal model (3) used in our testing procedure. This allows us to obtain more realistic data, and to assess the robustness of our method to deviations from the model.

We first generate a *q* × *p* peptide-protein membership matrix X. To do so, we assign to each protein a number of peptides drawn from a *P*(*q/p*) Poisson distribution. We make sure that each protein is assigned at least one peptide, and that all peptides are assigned to one protein. We then randomly select a subset of peptides to be shared across proteins (we show results for proportions 0%, 5%, 20% and 50%). For each of them, a Poisson draw with parameter 0.5 is used to determine the number of additional protein the peptide should connect to.

As explained in the introduction, the relationship between the measured peptide intensity and its actual abundance is peptide specific, so that the observed intensity of two protein-specific peptides from a same protein can be rather different (up to 3 orders of magnitude, according to Silva *and others* (2005)). To account for this fact, we multiply each positive entry of X by a draw from a 0.001 + *β*(1.1,3) distribution. In real cases, only the binary X membership matrix is known thanks to the protein sequence database: accordingly, the modified matrix is only used to simulate the intensities but only its binary counterpart is involved in our test.

We draw a 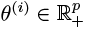 abundance vector for each of the *n* = *n*_1_ + n_2_ samples, in contrast to the model underlying our testing procedure where all samples under the same condition share the same *θ* parameter. More precisely, we draw one mean *θ* parameter for each condition from a log-normal distribution, then add independent normal perturbations around this mean for each sample (thresholding at 0 to make sure each 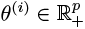). The mean and variance of protein-level log-normal distribution are chosen to fit classically observed datasets - that is peptide-level distributions that follow a Gaussian model centered on 23.5 and with a standard deviation of 2 on the log2 scale. The individual normal perturbations on the average abundance *θ_j_* of each protein *j* have mean 0 and standard deviation 0.005*θ*_*j*_. Under **H_*1*_**, the first 50 proteins are considered to be differentially abundant. The ratio between conditions is sampled from a normal distribution with mean R =15 and of standard deviation 0.05R.

Finally the log-abundance 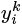 of peptide *k* in sample i is sampled from a normal distribution with mean log_2_ 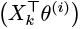 and variance 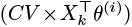 where X_k_ is the *k*-th row of *X* and where *CV* is a coefficient of variation that was set to 0.1 in our experiments.

The distribution from which we sample 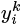 is therefore the log-normal (1), with a peptide-specific variance and a sample-specific (rather than condition specific) protein abundance, which is a bit more realistic. Importantly, the expectation of 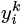 is not a linear combination of the parameter *θ*^(i)^, making our model (3) severely miss-specified. The R codes for our simulations is provided in the supplementary material.

### 4.2 Real datasets

The true set of differentially expressed proteins is generally not known in real data, making it difficult to compare differential analysis methods. We resort to using spike-in samples prepared according to the protocol of Ramus *and others* (2016). First, a lysate of yeast is split into *n* = 2*m*, so as to form an equal background for n samples. Then, a volume *V* of a mixture containing a series of precisely known human proteins is spiked in the first m samples so as to form the first biological condition. The other m samples receive a *R* × *V* volume of the same mixture so as to form the second biological condition with an abundance ratio *R*. As yeast and human proteins are different, any identified protein within each sample can uniquely be associated to its parent organism *(i.e.* yeast or human). During the relative quantification step, one should find that all and only the human proteins are differentially abundant.

We use the two datasets that are described in (Giai Gianetto *and others*, 2016), and which are available in the DAPARdata R package (Wieczorek *and others*, 2016). These two datasets contain 6 samples (*m* = 3) that have been prepared using the equimolar human protein mixture Sigma UPS1, including 48 human proteins. Their differential abundance ratios *R* are 2 and 2.5 for the first and second datasets, that are respectively referred to as Exp1_R2_pept and Exp1_R25_pept. Both datasets have been preprocessed with ProStaR software (Wieczorek and *others*, 2016), so as to make them compliant with the constraints of the reported experiments. First, all the peptides that corresponded to contaminant proteins or reversed proteins *(i.e.* peptides that were mistakenly identified as part of inexistent proteins) were removed, as well as those with more than one missing values out of three, in any of the two conditions. Second, the peptide intensities were normalized to account for replicate variability (within-condition median centering for Exp1_R25_pept, and global median centering for Exp1_R2_pept). Finally, the remaining missing values were imputed with an algorithm based on maximum likelihood estimation.

A limitation of this dataset is that it contains few shared peptides contrarily to real human samples, for a combination of reasons. First, the 48 human proteins available in the Sigma UPS1 equimolar mixture do not share peptides that are easily identified by mass spectrometry. Second, yeast is a simple organism that does not have a lot of homology sequences in its genome, so that there are very few shared peptides among its proteins. Finally, human and yeast proteins do not share many peptides, because they are rather different organisms. To cope with this issue, we artificially add shared peptides in these two real datasets, by merging pairs of peptides. Merging pairs of yeast peptides had no influence, as the resulting peptides (from non-differentially abundant yeast proteins) were even less prone to create error by introducing confusion between differentially and non-differentially abundant proteins. It was not possible to merge pairs of human peptides, because of their too limited number in the original datasets. We therefore resort to merging pairs of peptides where one is differentially abundant (human) and the other is not (yeast). To concretely implement such a merging procedure, a yeast peptide and a human one are randomly chosen, their identifications are merged, their abundances are summed, and their list of parent proteins are concatenated. This procedure can be repeated as many times as required to reach the desired amount of shared peptides. In Section 5, we report experiments with respectively 0, 120, 200 and 280 artificial shared peptides. These numbers are to be compared to the total numbers of human peptides in the datasets that are 211 (respectively 290) in Exp1_R2_pept (respectively Exp1_R25_pept).

### 4.3 Compared methods

We evaluate two versions of PEPA test, both using the likelihood ratio statistic (9): PEPA-ML relies on the ML estimator of the variance (7), and PEPA-MAP on the MAP estimator (8) of the variance with an inverse-gamma prior. Comparing these two procedures provides insights on the respective interests of the likelihood ratio test itself and the variance regularization. These two methods are compared to several other reference methods.

The only other peptide-based model that accounts for all shared peptides is AllP (Blein-Nicolas *and others*, 2012). However as explained in Section 2.2, it cannot cope with large scale datasets that we consider in our experiments and we compare to methods that only account for protein-specific peptides. The first one, referred to as PeptideModel, performs a two-sample t-test for each protein, where each group is formed by pooling all protein-specific peptides across all biological samples in one condition. This corresponds to a likelihood ratio test using model (3) without its peptide effect, and restricted to protein-specific peptides, so the performance increment between PeptideModel and PEPA-ML quantifies the interest of accounting for shared peptide in the context of this particular model. We also include the latest peptide-based method MSqRob from Goeminne *and others* (2016). Like PeptideModel, MSqRob relies on a linear model using the peptides as its sampling unit but introduces a few improvements: first, a ridge penalty on the estimated effects; second, an empirical Bayes estimator of the variance (both of which should help when few unique peptides are available); and finally, a robust loss function to deal with outliers. In the absence of shared peptides, it is therefore similar to PEPA-MAP but uses a different type of regularization of the variance, an additional regularization of the regression parameters and a robust loss.

We also consider two aggregation-based models: the first one, denoted AllSpec-SAM, performs the SAM-test proposed by Tusher and *others* (2001) (with automatic tuning of the fudge factor parameter, as discussed by Gianetto *and others* (2016)) over each protein summarized by the sum of intensities of all of its protein-specific peptides. The second one, denoted Top3Spec-TT, performs a t-test over each protein summarized by the sum of intensities of its 3 most abundant protein-specific peptides. AllSpec-SAM is the most accurate aggregation-based model, that is the best combination of aggregation and test, as discussed in Sections A and C of supplementary material. On the other hand, because of its simplicity, Top3Spec-TT remains one of the most used methods on proteomics platforms, and it provides baseline performances.

To compare the performances of the different methods, we construct precision-recall (PR) curves. In order to stabilize the results, the PR curves are averaged over 30 runs for simulated data. On real data, it is not possible to consider repetitions when no shared peptides are introduced (which makes the corresponding PR curve less smooth); however, when the random peptide merging procedure is called, 10 runs are considered each time to average the performances.

## 5 Results

In this section we present PR curves on simulated and real datasets, calibration curves and a runtime performance comparison of the evaluated methods.

### 5.1 Performances on simulated datasets

The performances on simulated datasets with 0%, 5%, 20% and 50% of shared peptides are displayed on Figure 1. Additional figures representing 1%, 10%, 33% and 67% of shared peptides are provided in Section D of supplementary material. From these results, one draws several conclusions:

**Figure 1:**
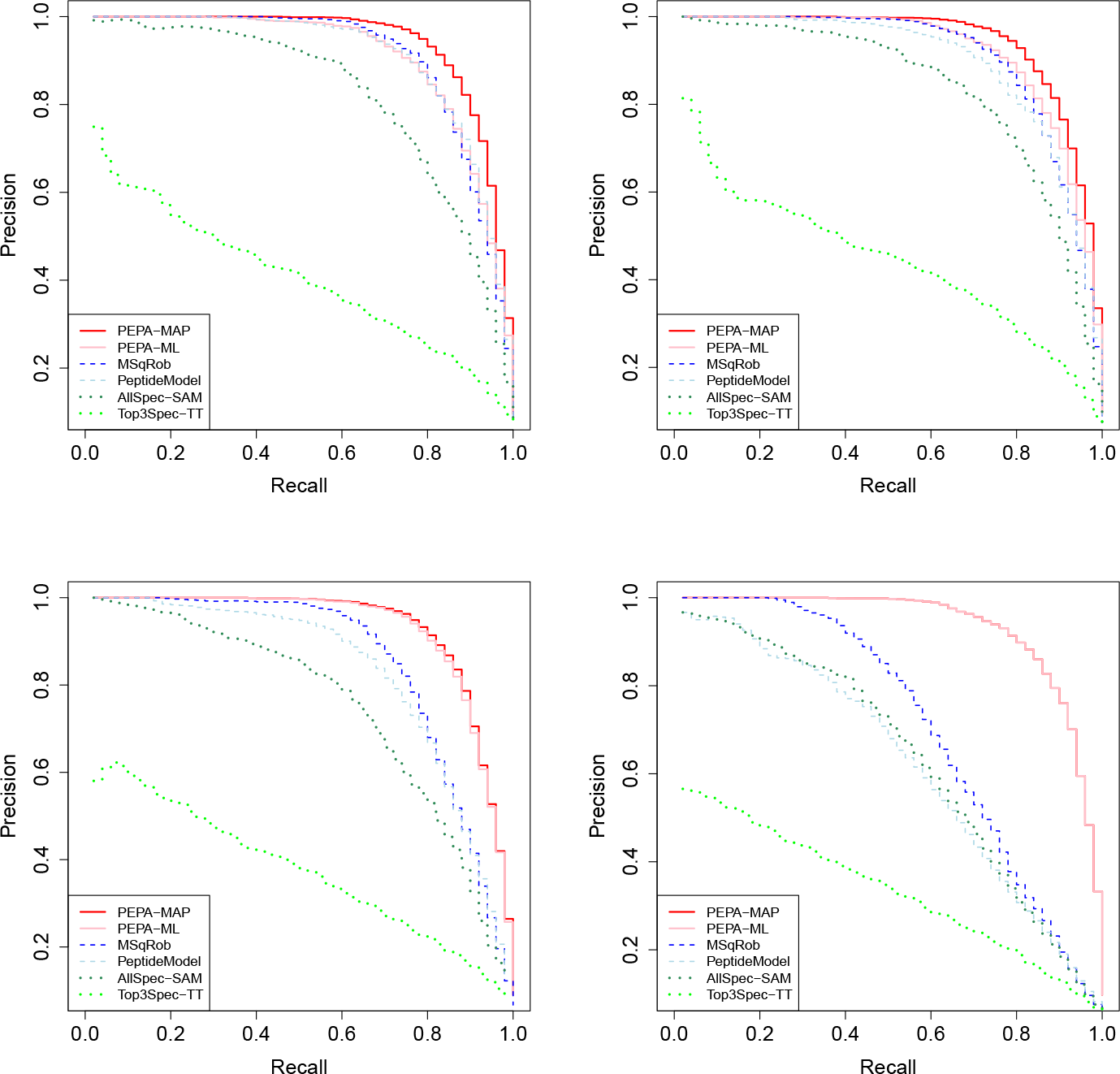
PR curve on simulated data with 0% (upper left), 5% (upper right), 20% (lower left) and 50% (lower right) of shared peptides.

#### Peptide-based dominate aggregation-based methods

In all settings, the baseline method Top3Spec-TT is by far the least accurate. Overall, as noticed by Goeminne *and others* (2015), aggregation-based methods (Top3Spec-TT and AllSpec-SAM, depicted by lighter or darker green dotted curves respectively) are less accurate than other methods, whether or not they exploit shared peptides.

#### Benefit of using shared peptides

In the absence of shared peptides, both PeptideModel and MSqRob are very similar to PEPA-ML and PEPA-MAP, respectively, as discussed in Section 4.3. Accordingly, the upper left panel of Figure 1 shows that both families perform comparably in this regime, whether they use shared peptides (PEPA-ML and PEPA-MAP in lighter or darker solid red curves respectively) or not (PeptideModel and MSqRob depicted by lighter or darker dashed blue curves respectively). As shared peptides are introduced, and as their number increases, the number of protein-specific peptides available mechanically decreases, affecting all methods which do not exploit shared peptides. On the other hand, the performances of our methods accounting for shared peptides are generally unaffected and clearly dominate all other methods as soon as a large enough proportion is reached (5 to 20%).

#### Benefit of regularization

The regularized versions of the compared methods (darker colors) always dominate their unregularized counterparts (lighter colors). This is true regardless of the proportion of shared peptides. As the number of shared peptides increases, the unregularized versions of methods which do not exploit these peptides (peptide-based PeptideModel and aggregation-based Top3Spec-TT) face a more severe dropout of their performances than the corresponding regularized methods (peptide-based MSqRob and aggregation-based AllSpec-SAM), suggesting that regularization helps more as fewer protein-specific peptides become available. We observe the opposite behavior for our methods accounting for shared peptides: the benefit of the regularization introduced in PEPA-MAP versus PEPA-ML decreases as the proportion of shared peptides increases. Indeed, as this proportion increases, our likelihood ratio test does not discard the shared peptides and additionally gains more peptides for each protein. Accordingly the pink curve of PEPA-ML actually improves as the proportion of shared peptides increases because its sample size also increases and regularization becomes less useful.

Overall, PEPA-MAP provides the best performances on simulated data, despite the strong misspecification of the data generating model with respect to the regression model.

### 5.2 Performances on real datasets

We build PR curves to compare all methods on Exp1_R2_pept (Figure 2) and Exp1_R25_pept datasets (Figure 3) with 0, 120, 200 and 280 shared peptides. Additional figures representing the cases with 40, 80, 160 and 240 shared peptides are provided in Section D of supplementary material. Exp1_R2_pept and Exp1_R25_pept originally contain 10722 (resp. 10601) peptides, among which 211 (resp. 290) come from human proteins, so that the proportion of shared peptides is rather small: for either datasets, the proportion of introduced shared peptides is smaller than 2.65% (to be compared with a proportion up to 50% of shared peptides as recalled in the introduction).

**Figure 2:**
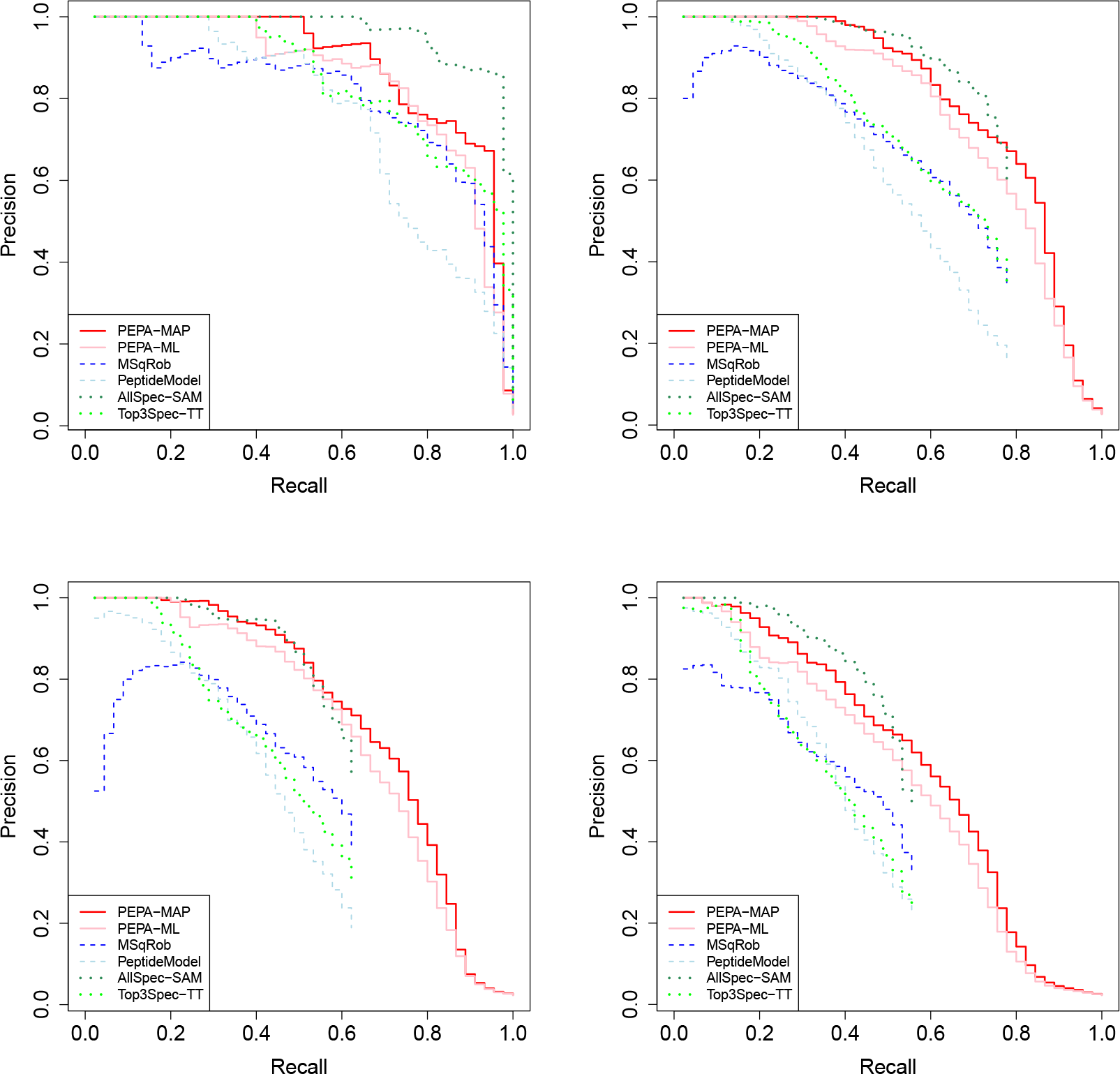
PR curve on Expl_R2_pept data with 0 (upper left), 120 (upper right), 200 (lower left) and 280 (lower right) artificially added shared peptides.

**Figure 3:**
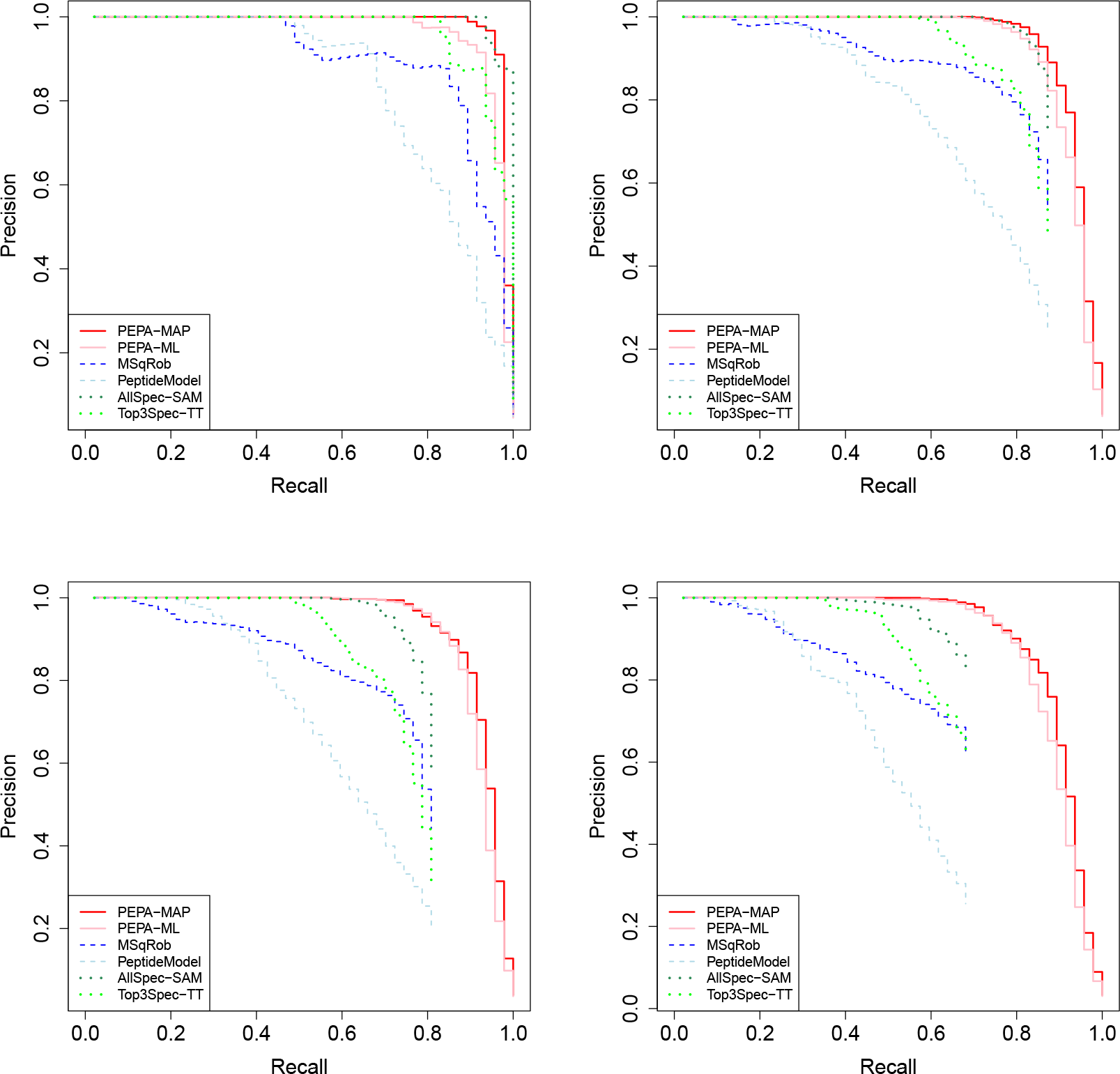
PR curve on Expl_R25_pept data with 0 (upper left), 120 (upper right), 200 (lower left) and 280 (lower right) artificially added shared peptides.

We notice a large difference of performances and of behavior between the two datasets. Expl_R25_pept derives from a series of LC-MS/MS experiments that did not undergo any malfunction, so that the data is of rather high quality. On Expl_R2_pept it is not possible to diagnose the source of the noise which could range from the cell acquisition process in the MS to the bioinformatic data processing, yet it is clear that on this dataset, the human UPS1 protein are more difficult to detect. Both datasets correspond to a scenario that can be faced on proteomics platforms.

#### Peptide-based methods do not always dominate aggregation-based methods

The domination of peptide-based over aggregation-based models that was clearly illustrated on simulated dataset does not hold on our real datasets. This is especially true on Expl_R2_pept, where AllSpec-SAM outperforms all other methods (including ours) when there are no shared peptides. In the presence of shared peptides, it is either competitive with or dominated by our method. All other methods (PeptideModel, Top3Spec-TT and MSqRob) are dominated by AllSpec-SAM, PEPA-ML and PEPA-MAP regardless of the number of shared peptides. A possible explanation is that when peptide-level intensity values are unreliable, the aggregation process somehow regularize the resulting protein-level intensity values. The same conclusions hold on the Expl_R25_pept dataset, yet with dimer magnitude: when the number of shared peptides increases, MSqRob keeps up but remains dominated by aggregation-based methods.

#### Benefit of shared peptides and regularization

PEPA-ML and PEPA-MAP outperform all other methods as soon as enough shared peptides are introduced. In all cases, the regularized methods AllSpec-SAM and PEPA-MAP outperform their unregularized counterparts Top3Spec-TT and PEPA-MAP. MSqRob dominates PeptideModel on Expl_R25_pept, but is outperformed for small recall values on Expl_R2_pept, suggesting that on this dataset MSqRob assigns the lowest p-values to a few non differentially abundant proteins.

#### Our methods handle proteins with no specific peptide

In both experiments, some proteins are lost by methods that only rely on protein-specific peptides because as the number of shared peptides increases, these proteins end up with only shared peptides. On Figures 2 and 3, this leads to the noticeable dropouts on the lower end of the curves for these methods. This illustrates an important practical issue: methods that only rely on protein-specific peptides are unable to deal with some proteins. Accounting for shared peptides like we suggest not only improves our ability to detect differentially abundant proteins among those that are handled by classical methods, but also increases the proteome coverage.

To conclude, these experiments show that both our method accounting for shared peptides and its regularized version improve our ability to detect differentially abundant proteins on proteomics datasets. The more shared peptides, the more important the increment of performances, but even on datasets with no shared peptides the methods remain accurate and never strongly underperforms. Finally, the regularized version always performs better than the other, making PEPA-MAP a safe choice.

### 5.3 *p*-value calibration

The last point to evaluate is the quality of the calibration of the *p*-values provided by our tests. In particular, it is necessary to check that the correction we introduced in Proposition 2 for our PEPA-MAP statistic leads to correct asymptotic levels using a χ^2^ distribution like with the PEPA-ML statistic. To visually assess this point, we compare the expected and actual test levels for the methods evaluated in Sections 5.1 and 5.2 except for Top3Spec-TT, to avoid cluttering our graphs. In addition to PEPA-MAP we include a corrected version PEPA-MAP-RW. RW stands for reweighted: all the regularized likelihood ratio statistics are multiplied by 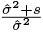 as suggested by Proposition 2. We compute the mean square residuals of our model for each protein and average these estimates across proteins to obtain 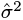. PEPA-MAP-RW would behave exactly like PEPA-MAP in the PR curves of Sections 5.1 and 5.2 as multiplying the test statistic of all proteins by the same weight does not affect their order. The p-values for PEPA-ML, PEPA-MAP and PEPA-MAP-RW are computed by comparing the corresponding statistics to the quantiles of a 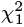 distribution.

Figure 4 is a (log-log) plot of the empirical proportion of false positives obtained as a function of the *p*-value threshold, *i.e.,* the proportion of nondifferentially abundant proteins (y-axis) which are assigned a p-value lower than the threshold (x-axis). If a test is correctly calibrated, a proportion *α* of nondifferentially abundant proteins has p-value lower than *α* for all *α ∈* [0, l] and its calibration plot is the *y* = *x* axis. Additional figures representing the various calibration plots we obtained during the experiments are provided in Section D of supplementary material.

**Figure 4:**
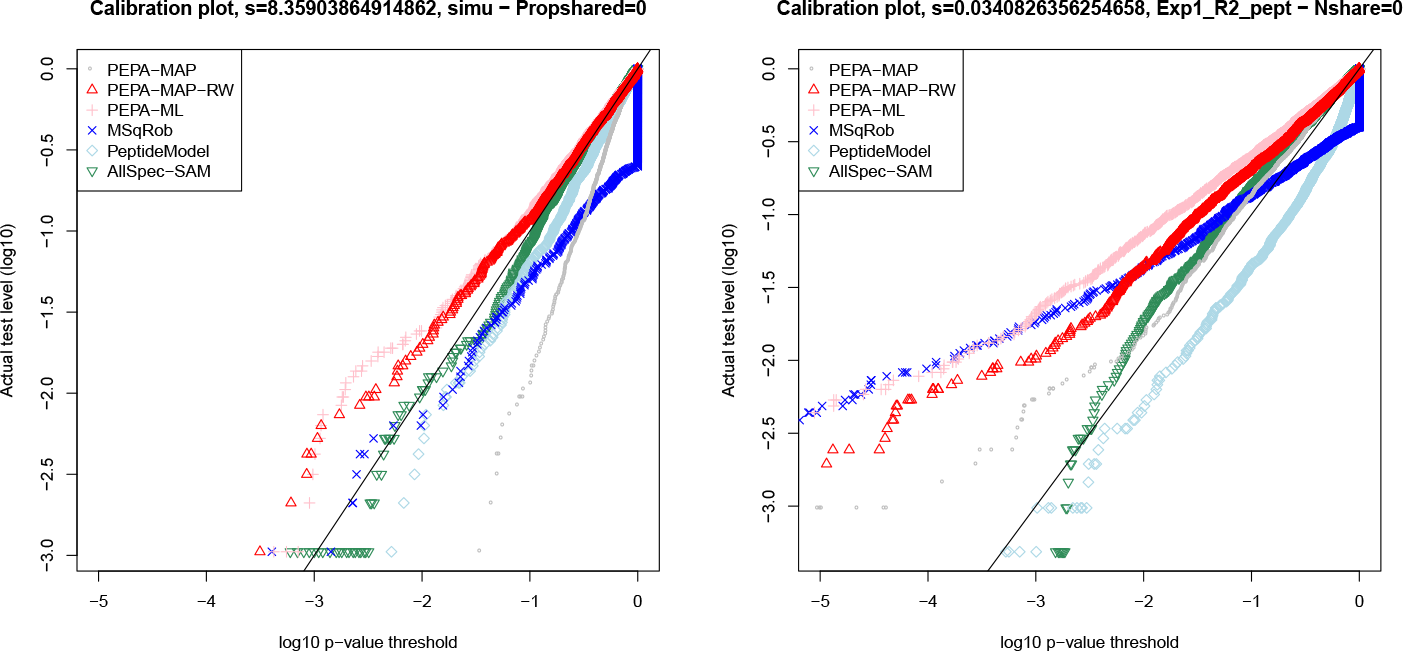
Calibration plots on simulated (left) and Expl_R2_pept (right) data for all compared testing procedures.

#### Calibration of PEPA-ML

The left panel of Figure 4 shows the plots obtained on simulated data from Section 4.1 with no shared peptide. All methods except PEPA-MAP are reasonably well calibrated. The small deviation observed for PEPA-ML can be explained by a combination of two factors. First, the χ^2^ distribution of PEPA-ML is an asymptotic result, and we only have *n*_1_= *n*_2_ =3 observations for each group in this case. Second, the null hypothesis that we are testing is *θ* = *θ*^’^, *i.e.,* that no protein is differentially abundant. This model is misspecified as soon as the protein *j* that we are testing is in the same connected component as a differentially abundant protein *j*^ș^, i.e. *θ*_j^’^_ ≠ 0, even if indeed *θ*_*j*_ = 0. This may happen in our simulations, as our dataset contains 50 differentially abundant proteins. We observe nonetheless that using the same simulation setting with 10 samples per group instead of 3 leads to a perfectly calibrated PEPA-ML (not shown), suggesting that the main issue is the low sample size. The right panel of Figure 4 shows the plots obtained on the Exp1_R2_pept data. PEPA-ML is more severely decalibrated, leading to a false positive rate of 7.3% when thresholding at 0.01 when AllSpec-SAM leads to a 1.9% rate, PeptideModel to 0.4% and MSqRob to 4.3%. The deviation is likely caused by the low sample size, as the number of differentially abundant proteins in the dataset is very similar to the one used in our simulation - where we recover a correct calibration by increasing the sample size.

#### Calibration of PEPA-MAP

As predicted by Proposition 2, the PEPA-MAP statistics are not 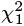 distributed under **H**_0_ which is illustrated by the strong deviation of the grey curve from the *y* = *x* axis - the selected regularization parameter s is large. Weighting our regularized statistics by the factor obtained in Proposition 2 leads to a test with similar calibration as our unregularized PEPA-ML, i.e., whose small deviation from the correct level can be explained by the low sample size. On the Exp1_R2_pept data the deviation of PEPA-MAP is milder because the selected s is smaller. It actually leads to more accurate levels than PEPA-ML (1.4% false positive rate when thresholding the p-values at 0.01) by partially compensating the deviation incurred by PEPA-ML (because of small sample size) in the opposite direction. This is of course artefactual and should not be considered a good property as there is no guarantee the same phenomenon will systematically happen on new data. The weighting scheme mostly corrects the deviation of the levels of PEPA-MAP from those of PEPA-ML, leading to a 4.4% false positive rate when thresholding the p-values at 0.01. The remaining difference is probably caused by the poor quality of our estimate of 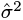, and the fact that the data may not be well represented by i.i.d samples from a distribution with a common variance.

Additional calibration plots with varying number of artificially added peptides for Expl_R2_pept and for Expl_R25_pept are displayed in Section D of supplementary material.

### 5.4 Runtime evaluation

Table 1 shows the average runtime of all evaluated methods across five runs of the synthetic experiment described in Section 4.1. All coefficients of variation are below 10%. We only show a single result denoted as PEPA for PEPA-ML and PEPA-MAP as the marginal cost of adding a fudge factor is small. For PEPA, we also show the average time spent at computing the test statistic. The rest of the time is used to identify the connected component of the peptide-protein graph, and is unaffected by the speedup obtained in Proposition 1. We do not show the execution time of PEPA without the speedup but each experiment takes more than ten hours.

**Table 1:** Average execution time in seconds across five runs for the evaluated methods on simulated data with 0%, 5%, 20% and 50% of shared peptide.

| Shared peptides | Top3Spec-TT | AllSpec-SAM | PEPA (test) | MSqRob | PeptideModel |
| --- | --- | --- | --- | --- | --- |
| 0 | 1.67 | 0.44 | 56.84 (1.85) | 403.36 | 0.63 |
| 0.05 | 1.55 | 0.49 | 60.16 (1.76) | 494.20 | 0.61 |
| 0.2 | 1.43 | 0.55 | 64.77 (5.61) | 660.61 | 0.57 |
| 0.5 | 1.13 | 0.49 | 67.74 (5.93) | 992.35 | 0.50 |

Most methods which do not use shared peptides have a runtime close to one second: computing their statistic only involves simple operations such as averages over small numbers of observations. MSqRob however is two orders of magnitude slower as it involves more observations and an iterated reweighted least square procedure, but it remains fast enough to be applicable to proteomics datasets with hundreds of proteins and thousands of peptides. Our methods run in one minute, most of which is spent computing the connected components of the peptide-protein graph. The computation of our test statistic using Proposition l takes less than 10 seconds, even though it requires the residuals of a linear regression problem with an *nq* × (*p* + *q)* design matrix.

## 6 Discussion and conclusions

We have proposed a linear model that accounts for shared peptides in relative quantification proteomic experiments based on mass-spectrometry analysis of peptides. This model can be used to build likelihood ratio tests relying either on the maximum likelihood or maximum a posteriori estimator of its variance parameter. We have also introduced a faster way to compute the test statistic, making it amenable to datasets with thousands of peptides and proteins. The faster form relies on the fact that the likelihood ratio statistics only requires the regression residuals as opposed to estimates of the regression parameters, in particular of protein abundances. Using our model to estimate abundances - a task which is out of the scope of this paper and which we did not include in our experiments - would require to actually estimate the regression parameters, *e.g.* using explicit formulas for 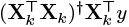 as suggested in Section 3.2.

Experiments on simulated and spike-in data confirm that the proposed tests have a clear advantage against existing methods to detect differentially abundant peptides in the presence of shared peptides. In the absence of shared peptides or when very few of them are present, our tests behave like existing methods, suggesting they can be safely used in all cases. We have also shown that asymptotic levels could be obtained when using the maximum a posteriori estimator instead of the maximum likelihood, providing asymptotic levels for this version of our likelihood ratio test - which systematically outperforms the maximum likelihood version in our experiments. An implementation of our tests is available via the `pepa.test` function of the Bioconductor package `DAPAR`.

Our work could be extended in several ways. First the experimental design of some proteomic analyses may be more complex than the one accounted for in our evaluations. The fast version of our statistic introduced in Proposition 1 is derived for a model with a protein and peptide fixed effect only, as opposed to *e.g.* technical replicates. Proposition 1 could be generalized to more grouping factors with fixed effects. Alternatively, or for random effects, one can use model (3) with additional factors and (slower) out of the box implementations of mixed model to compute the likelihood ratio statistic. Another possible extension regards the misspecification described in Section 5.3: using a Wald test instead of likelihood ratio test, one could simply estimate all protein abundances jointly instead of relying on models in which all proteins but one are differentially abundant. Wald tests however would not benefit from the acceleration allowed by Proposition 1 as they require parameter estimates as opposed to just likelihoods.

## Supplementary Materials

The reader is referred to the Supplementary Materials for additional comparisons between methods, additional PR and calibration plots as well as for the R code used in our experiments.

## Acknowledgments

The authors thank Samuel Wieczorek for his help in integrating the PEPA test to the DAPAR package and Zaïd Harchaoui for insightful discussions.

